# Multiethnic catalog of structural variants and their translational impact for disease phenotypes across 19,652 genomes

**DOI:** 10.1101/2020.05.02.074096

**Authors:** Fritz J. Sedlazeck, Bing Yu, Adam J. Mansfield, Han Chen, Olga Krasheninina, Adrienne Tin, Qibin Qi, Samantha Zarate, Joshua L. Traynelis, Vipin Menon, BCM HGSC Sequencing Lab, Jianhong Hu, Harsha Doddapaneni, Ginger A. Metcalf, Josef Coresh, Robert C. Kaplan, Donna M. Muzny, Goo Jun, Richard A. Gibbs, William J. Salerno, Eric Boerwinkle

**Affiliations:** Human Genome Sequencing Center, Baylor College of Medicine, Houston, TX; School of Public Health, University of Texas Health Science Center at Houston, Houston, TX; Bloomberg School of Public Health, Johns Hopkins University, Baltimore, MD; Department of Epidemiology and Population Health, Albert Einstein College of Medicine, Bronx, NY; DNAnexus, Mountain View, CA; Division of Public Health Sciences, Fred Hutchinson Cancer Research Center, Seattle, WA

**Author notes:** These authors contributed equally to this work (Fritz J. Sedlazeck, Bing Yu). see Supplement for list of collaborators (BCM HGSC Sequencing Lab). Corresponding author: Eric Boerwinkle, Human Genetics Center, University of Texas Health Science Center at Houston, 1200 Herman Pressler, Suite W-102, Houston, TX 77030.

## Abstract

Genome sequencing at population scale provides unprecedented access to the genetic foundations of human phenotypic diversity, but genotype-phenotype association analyses limited to small variants have failed to comprehensively characterize the genetic architecture of human health and disease because they ignore structural variants (SVs) known to contribute to phenotypic variation and pathogenic conditions^1–3^. Here we demonstrate the significance of SVs when assessing genotype-phenotype associations and the importance of ethnic diversity in study design by analyzing SVs across 19,652 individuals and the translational impact on 4,156 aptamer-based proteomic measurements across 4,021 multi-ethnic samples. The majority of 304,533 SVs detected are rare, although we identified 2,336 protein-coding genes impacted by common SVs.\

We identified 64 significant SV-protein associations that comprise 36 cis- and 28 trans-acting relationships, and 21 distinct SV regions overlapped with genome-wide association study loci. These findings represent a more comprehensive mapping of regulatory and translational endophenotypes underlying health and disease.

## Main

Targeted sequencing has demonstrated the importance of SVs in studies of highly mutated genomes (e.g. cancer) or rare Mendelian conditions attributable to large copy number events^2,3^. Sudmant, et al. ^4^ demonstrated the characteristics of SVs in 2,504 multiethnic human genomes, and despite the limited sample size and relatively low coverage, the data from this study have proven invaluable for biomedical research and translation. Other studies^5,6^ have described the relationship between SVs and gene expression, though on a small number of samples (n=147). These multi-sample, multi-omic SV studies have occurred in parallel with refinement of SV detection, genotyping and annotation algorithms, the construction of high-confidence SV “truth” sets ^7^ and ready access to high-throughput computing, all required for population-scale SV analysis.

We analyzed whole genome sequence data for the occurrence and characteristics of rare and common SVs in 19,652 European-American, Hispanic/Latino- and African-American genomes (see **Methods** and **Supplementary Section 1-2**). SVs were detected with the Parliament2 multimethod consensus SV framework^8^ and joint-genotyped with muCNV, resulting in 304,533 SVs. SV genotypes were then related to comprehensive aptamer-based plasma proteomic data (4,156 proteins) from 4,021 individuals as a metric of the impact of SVs on genome function. The SV data are available on dbVar (nstd160) and as a downloadable genome browser track.

The majority of the detected SVs were deletions, mostly in the 20-50 bp and 1-10 kbp size ranges, while duplications, the second most prominent SV type, were mainly observed in the 1-10 kbp size range. **Figures 1A and 1B** summarize the frequency of the observed deletions across size regimes and within each ethnicity: African-American (N=2,584), Hispanic/Latino-American (N=8,099) and European-American (N=8,969). These ethnicities segregate as expected in Principal Component Analysis (PCA) of the SVs relative to single-nucleotide variant PCA (**Figures 1C and 1D**). African-Americans have the highest overall MAF (median 0.058%) compared to European-Americans and Hispanic/Latino-Americans, with median MAFs of 0.017% and 0.012% respectively. Even excluding singleton SVs this difference persists, with median MAFs of 0.077%, 0.033% and 0.025% for African-Americans, European-Americans and Hispanic/Latino-Americans, respectively. Clustered breakpoints in common SVs (MAF >0.05 in at least one ethnicity) identify regional misalignments that may impact SNP calling and clinical interpretation. We identified 31.57 Mbp (African-American), 24.92 Mbp (Hispanic/Latino-American) and 25.81 Mbp (European-American) that are within 1 kbp of two or more breakpoints of common SVs (**see Supplementary Section 2**). When analyzing FST in10kbp sliding windows, we observe an average FST of 0.033 (sd:0.03) that is likely heavily influenced by admixture of Hispanic/Latino-Americans. A pairwise FST comparison revealed that African-vs. European-Americans (average FST:0.09, sd:0.11) have the highest, followed by African- vs. Hispanic/Latino-Americans (average FST:0.05, sd: 0.07) and then European vs. Hispanic/Latino-Americans have the lowest (avg. FST: 0.01, sd: 0.01) (see **Supplement Table 12**).

**Figure 1:**
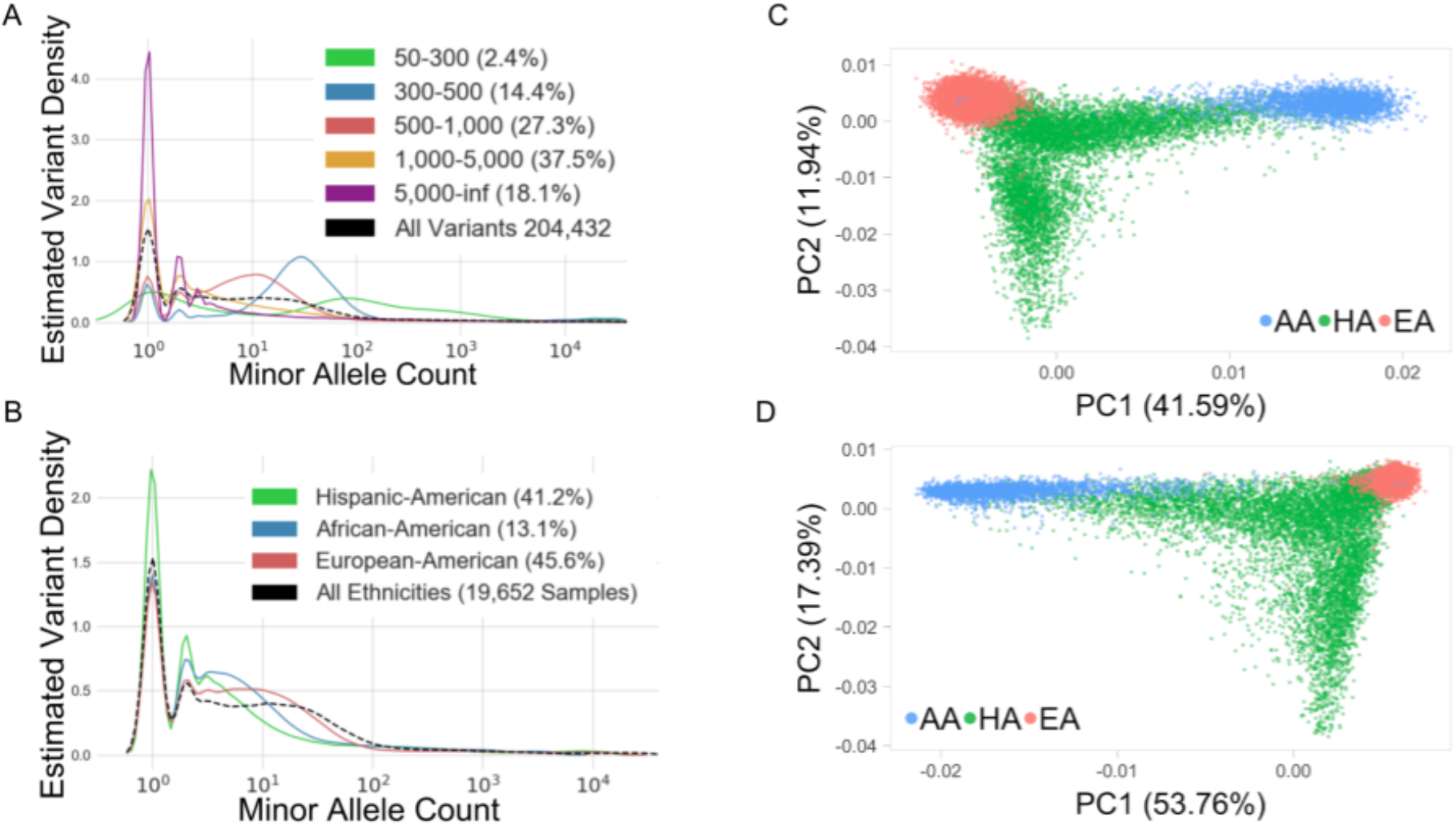
Population SV summary. SV minor allele count spectra for deletions by event size (A) and ethnicity (B) across 19,652 CVD samples. (C) Principal components 1 and 2 for common (MAF >0.1) deletions colored by ethnicity. (D) Principal components 1 and 2 for common (MAF >0.1 SNVs on chromosome 1.

Applying similar methods, we also identified 308,867 SVs in ten non-human primate (chimpanzee and gorilla) samples to annotate the human SVs with their likely ancestral alleles and characterize differences between human and non-human primates. Compared to the catalog of human SVs, 1,013 events were directly overlapping (+/-100 bp) and 1,240 were overlapping within +/-1 kbp. The vast majority (93.02%) of the overlapping SV are common (AF>0.05) and are deletions (77.82%) in both human and in non-human primates. However, we did observe 266 (21.45%) SVs with different SV types (eg. Deletions vs. duplications) between species likely identifying shared regions of genome instability (see **Supplementary Section 3**).

Of the 304,533 SVs, 118,989 (39.07%) overlap with 13,398 (66.17%) annotated protein-coding genes (including introns). The majority of these SVs are rare (MAF <0.05 in each ethnicity). We identified 2,336 genes impacted by common SVs (MAF >0.05 in at least one ethnicity). Limiting this analysis to African-Americans, this number is 2,172, underscoring the benefits of increased diversity in genotype-phenotype association studies. **Figure 2A** shows the set of shared genes impacted by common SVs among ethnic groups across multiple MAF thresholds. The majority of SVs (51.63%) are present within each of the three populations and represent common SVs in the broader human population and are likely ancient in origin. Considering the 1,649 genes having SVs found in all three populations (see **Supplementary Section 4**), there is significant GO term enrichment for antigen binding (GO:0003823 FDR q-value: 1.36E-7), leukocyte migration (GO:0050900 FDR q-value: 3.19E-6) and immune system process (GO:0002376, FDR q-value: 1.67E-4). To identify ethnic-specific gene-associated SVs we filtered for MAF >0.05 for African-, Hispanic/Latino- or European-Americans while requiring that the events also have an MAF <0.05 in the other two populations. **Figure 2B** shows the overlap of these genes that are impacted by ethnic-specific SVs. Only three genes (*GUCY1A2, PRH1-PRR4, PRR4*) are impacted by at least one distinct ethnic-specific SV for each of the three populations. *GUCY1A2* is impacted by three different non-overlapping deletions targeting different regions that are ethnic-specific. *PRR4* and *PRH1-PRR4* show ethnic-specific deletions for African- and European-Americans, but two slightly overlapping (by 33 kbp) duplications common in Hispanic/Latino-Americans, suggesting polymorphic *PRR4* variation possibly increasing that gene’s expression in the Hispanic/Latino population^9^. When assessing the molecular function of these genes impacted by ethnicity specific SVs, we observed that the majority of the genes are associated with binding (GO:0005488) and catalytic activity (GO:0003824).

**Figure 2:**
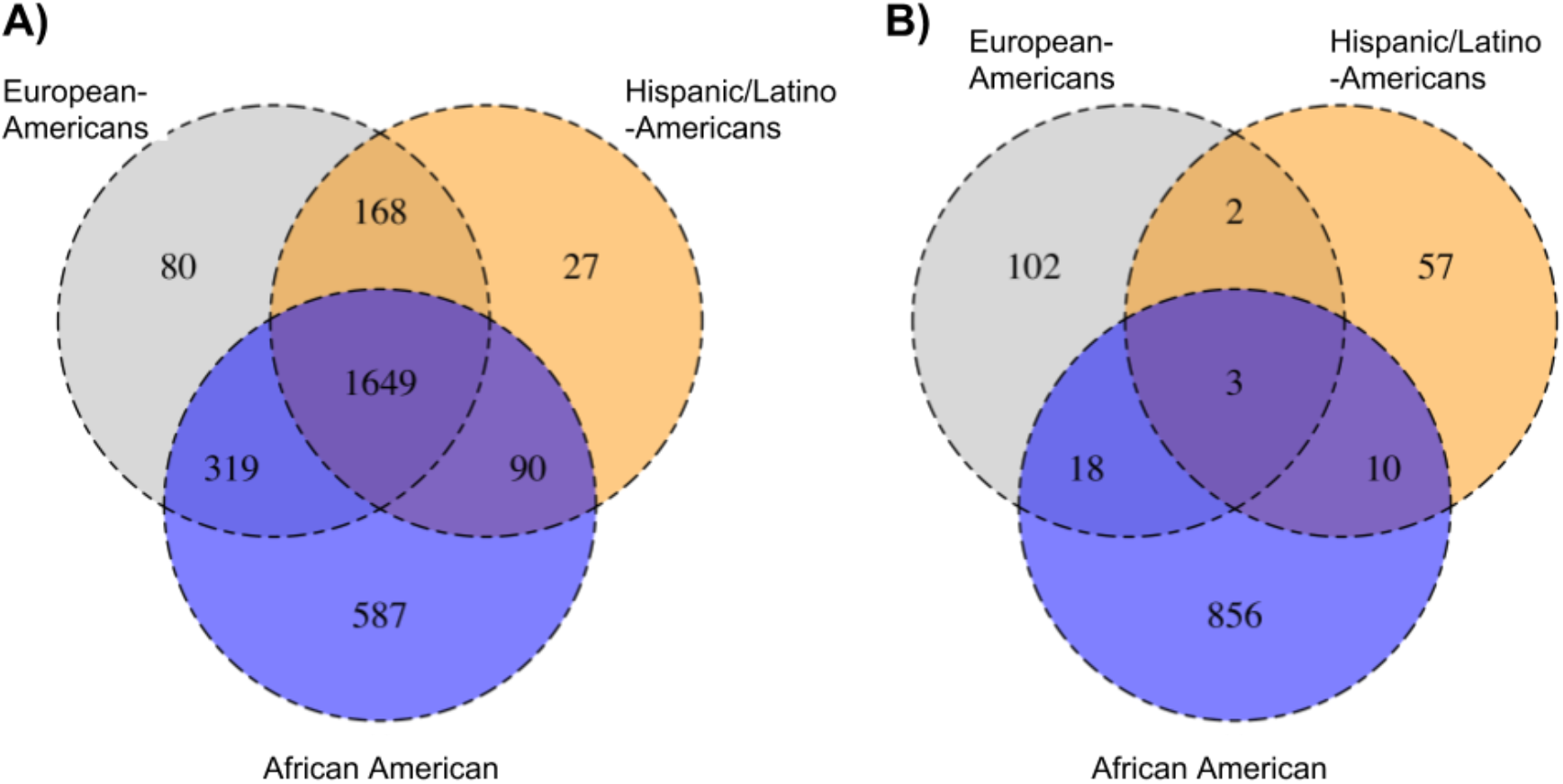
Overlap of genes influenced by SVs. A: Genes that are overlapping SVs common in specific ethnicities: European- (gray), Hispanic/Latino- (yellow) and African-Americans (blue). B: Genes that are overlapping by SVs common in one ethnic population (eg. MAF >0.05), but rare (e.g. MAF <0.05) in the other ethnic population. The three-ethnicity overlap indicates multiple genes that are impacted by SVs specific to certain populations. Supplement Figure 4 shows the impact across genes for multiple MAF thresholds.

One advantage of genome sequencing over that limited to the exome is the ability to interrogate regulatory regions outside the protein-coding portion of the genome. Therefore, we next identified known regulatory elements impacted by SVs. For SVs common in any one ethnicity (MAF >0.05), we identified 9,445 (10.25%) different CTCF binding sites impacted by SVs, followed by 8,668 (6.70%) open chromatin regions, 4,241 (9.28%) promoter flanking regions and 2,663 (7.31%) enhancers. These regulatory elements are more often impacted by deletions (59.13%) than duplications (38.60%) or inversions (2.27%). For ethnic-specific SVs (MAF >0.05 in only one ethnicity), we observed 2,154 different regulatory elements impacted in African-Americans while only 236 and 387 for European- and Hispanic/Latino-Americans, respectively. Considering different SVs but within the same gene, the highest degree of overlap was between African-Americans with European-Americans (18 genes) and African-Americans and Hispanic/Latino-Americans (10 genes) rather than all three ethnicities (3 genes) (**Figure 2**). For African-American-specific SVs, deletions (565) more often impacted regulatory elements than duplications (312), whereas in European-Americans this trend seems to be the opposite (DEL: 57, DUP: 83) (see **Supplementary Section 4**).

These analyses characterize structural variation across the human genome and describe the potential impact of these SVs based on ethnicity and overlap with genes and regulatory elements. However, the pathways by which SVs influence health and disease are varied, including transcription as analyzed elsewhere^5^ and translation as described here. We selected the proteome as one over-arching metric to assess the impact of SVs on biologic function. The effects of 60,433 SVs were analyzed on 4,156 aptamer-based plasma protein levels^10^ that passed QC (**see Methods**) among 4,021 European- and African-Americans from the Atherosclerosis Risk in Communities (ARIC) study. We identified and replicated 64 unique SV-protein associations, comprising 44 deletions, 16 duplications and 4 inversions (**Figure 3**), among which four of the associations were reported by the previous SV-transcription study^5^. The 64 identified SVs mapped to 41 distinct regions, and 21 of these regions link to 111 disease-related traits reported in genome-wide association studies (**Supplemental Table 6-7**). The median phenotypic variance for the proteome explained by the 64 SV was 16.1% for European-Americans and 11.3% for African-Americans. Thirteen of the 64 SVs overlapped the gene coding for the protein targeted by the corresponding aptamer. For example, a 22,300 bp deletion in *GSTM1* is associated with decreased levels of Glutathione S-transferase Mu 1, a protein encoded by *GSTM1* that detoxifies electrophilic compounds, such as carcinogens and products of oxidative stress^11,12^. Similarly, a 4,200-bp duplication in *CR1* is associated with increased levels of Complement receptor type 1, a protein encoded by *CR1* that plays a major role in immune complex clearance^13^. Among the remaining 51 SVs, 23 have effects on proteins that are encoded by nearby genes (i.e. *cis*-acting), and 28 have effects on proteins that are encoded by distant genes, defined as different chromosomes (i.e. *trans*-acting). One example of SV complexity reflecting translational complexity is a variant genotyped as both deletion and duplication events, showing cis- and transacting effects and changes in protein level that are dependent on the copy number. A 19,300-bp deletion and a 17,400-bp duplication in *SIGLEC14* (19q13.41) is significantly associated with decreased levels of Cathepsin L1, a major controlling element of neutrophil elastase activity, which is encoded by *CTSL* on chromosome 9. *SIGLEC14* is adjacent to *SIGLEC5*, which undergoes partial gene conversions with *SIGLEC14* in primates^14^. These deletions and duplications are also associated with levels of the *SIGLEC14* and *SIGLEC5* proteins. Neutrophil elastase is well known for its antibacterial activity^15^, and *SIGLEC* protein levels regulate the functions of cells in the innate and adaptive immune systems^16^. A previous study suggested that the deletion of *SIGLEC14* may play a role in bacterial infection^17^. Of note, there are 16 associations where deletions are associated with increased protein levels. For example, a 1,500-bp deletion in 10q11.22 is associated with increased levels of beta-microseminoprotein, a biomarker of prostate cancer encoded by *MSMB*. The deletion is located approximately 30 kbp upstream of transcriptional start site, immediately proximal to the enhancer region for *MSMB*. We speculate this deletion promotes transcription of *MSMB*, which may lead to a protective effect on prostate cancer as lower *MSMB* expression and beta-microseminoprotein levels are associated with increased risk of prostate cancer^18,19^. We investigated the association between protein levels and deletion size and observed that large deletions were enriched in the set of significant and replicated protein associations (p<0.001). For those detected SV regions, we included both single nucleotide variants (SNV) and SV into an integrated analysis of protein levels. After conditioning on genome-wide significant SNVs, SV-protein associations were attenuated, affecting more *trans* acting than *cis* acting associations. Of note, 16 out of 58 (27.6%) and 11 out of 33 (33.3%) associations in the discovery set remained significant in European- and African-Americans, respectively (**Supplemental Table 8-9**). On a per-variant basis, SVs had an average of 1.78 fold greater effect size than SNVs, suggesting a predominant role of SV in the detected region (**Supplemental Table 10-11**).

**Figure 3:**
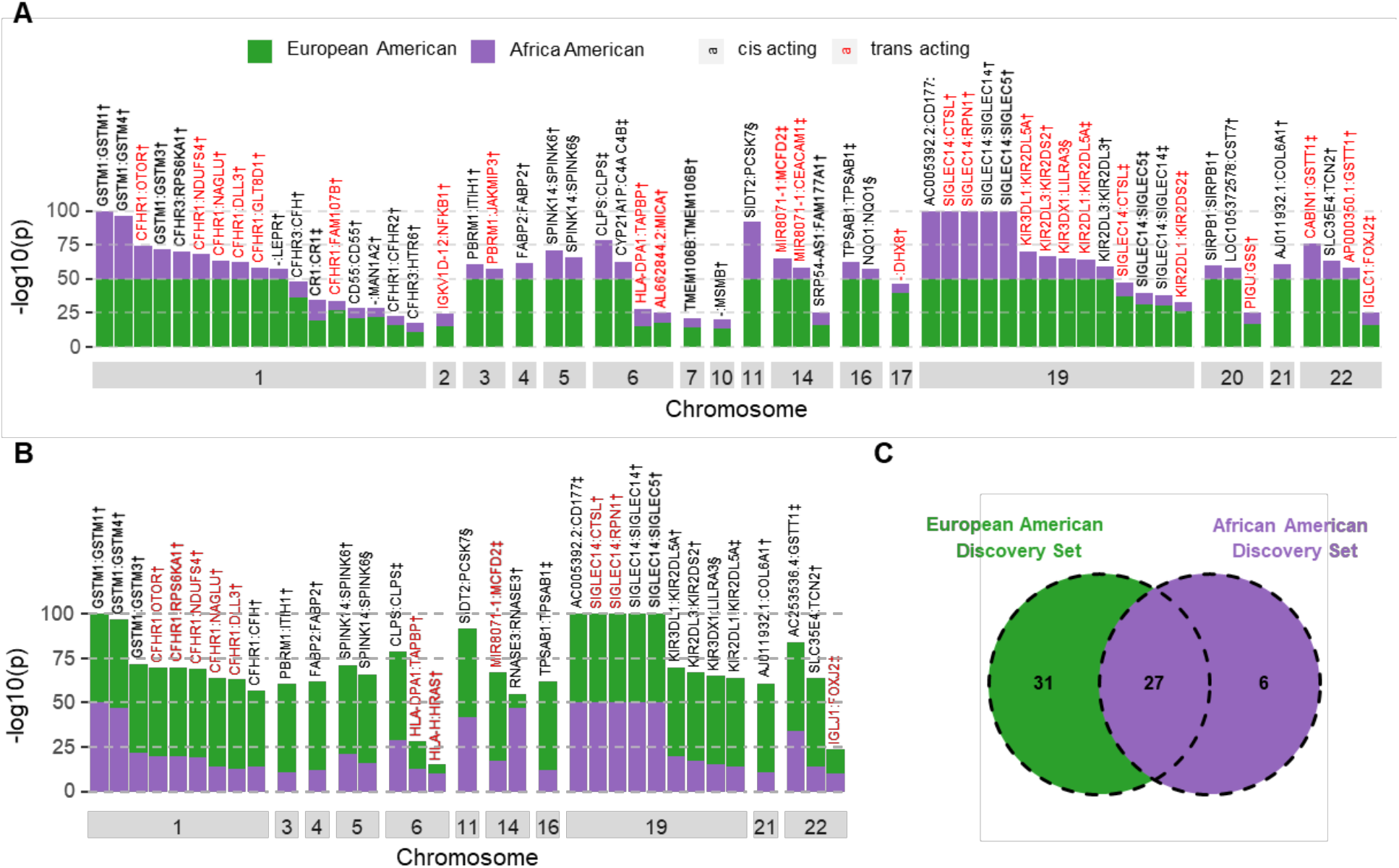
A) Bar plot of –log10 p-values for fifty-eight structural variation related proteins in European Americans (discovery) and African Americans (replication) by chromosome; B) bar plot of –log10 p-values for thirty-three structural variation related proteins in African Americans (discovery) and European Americans (replication) by chromosome; and C) the number of identified structure variation-protein intersections between European American discovery set and African American discovery set. The length of each color bar represents —log10 p-value in the corresponding study sample set, which is truncated at 50 if p-value < 1e-50. Bars are labelled as structural variant overlapping gene: protein coding gene. † indicates deletion event, ‡ indicates duplication event and § indicates inversion event. Structural variation-protein pairs in red indicate *trans*-acting effects.

Large-scale genome sequence generation from programs such as TOPMed, Genomics England, All of Us and the Centers for Common Disease Genomics is outpacing the community’s ability to fully leverage whole-genome information for novel discovery, diagnostics and genomic medicine. The results presented here demonstrate the importance of ethnically inclusive study design and the incorporation of SVs for comprehensive large-scale genomic studies. Harmonized SV catalogs such as those presented here serve not only as a resource for genome annotation, improved variant interpretation and expanded disease-gene discovery, but also as a model for further at-scale multi-omics data integration. As the number of genome sequences reaches and surpasses millions, technical challenges will continue to require innovative scalable logistics, compute and quality control solutions. However, the limits of current short-read DNA sequencing data will motivate the field toward long-range sequencing and other omics data, just as the limits of capture and limited exome experiments catalyzed it to sequence deep genomes of ever-larger cohorts. We anticipate that judicious use of long-range sequencing in the near term will improve the utility of existing short-read genome data rather than supplant it entirely.

## Methods

### Study Population

The Atherosclerosis Risk in Communities Study (ARIC) is an ongoing biracial cohort designed for cardiovascular research, as described in detail previously^20^. In brief, a total of 15,792 individuals, predominantly African- and European-Americans, participated in visit 1 during 1987-1989, with three additional triennial follow-up visits, a fifth visit in 2011-2013 and a sixth visit in 2016-2017. The Hispanic Community Health Study / Study of Latinos (HCHS/SOL) is a community-based cohort study of Hispanics/Latinos designed to examine risk and protective factors for chronic diseases, as published previously^21,22^. In brief, 16,415 self-identified Hispanic/Latino Americans of Mexican, Puerto Rican, Dominican, Cuban, or Central and South American origin were recruited from four field center. The HCHS/SOL participants underwent their visit 1 during 2008-2011 and visit 2 in 2015-2017.

### SVs calling and annotation

All 19,853 blood-derived DNA samples were sequenced on the Illumina HiSeq X platform (150 x 150 bp paired-end reads) to an average depth of >30x (see **Supplementary Section 1**). FASTQs were processed via the NIH-compliant GRCh38 protocol ^23^, generating per sample CRAM files for SV discovery and genotyping. Single sample SV discovery was performed with Parliament2^8^, a consensus SV framework that integrates multiple SV methods to identify high-quality SV events.

Parliament2’s consensus strategy is optimized against the NIST GIAB HG002 high-confidence SV (deletion F-measure=81.73) and provides quality scores informed by the size, type and method-support combination for each event. For this callset, Parliament2 (using default parameter) was run on each sample via a single Docker image across multiple cloud platforms (DNAnexus, AWS, Google Cloud Platform), generating a single consensus VCF for each sample. Sample VCFs were merged across all samples with SURVIVOR^9^ (with 1kbp distance and typesave parameters), resulting in a non-redundant set of candidate SVs (4,561,050 deletions, 3,239,510 duplications, 132,773 insertions, 2,359,206 inversions). These candidate SVs were then jointly-genotyped across the 19,853 CRAM files with muCNV (see **Supplementary Section 2**), yielding a raw set of 464,025 jointly genotyped SVs (258,667 deletions, 79,910 duplications, 125,448 inversions).

These genotyped SVs were then quality controlled to remove outlier variants and samples. Clusters of overlapping SVs (reciprocal overlap >0.5, non-reference genotype concordance >0.8) were trimmed to the single event of the highest quality. Artefactual duplication calls, defined as duplications with MAF >0.5 and no support from read-pair mappings or peripheral read depth, were excluded from the final data set. Using the filtered genotypes, variants were counted per sample and 171 outlier samples (>9,000 SVs each) were removed from the callset. The final SV callset comprises 19,652 samples (2,584 African American, 8,969 European American, and 8,099 Hispanic/Latino American) and 304,533 SVs (204,432 deletions, 44,060 duplications, and 56,041 inversions) described in **Figure 1 and Supplementary Table 2**.

Each candidate SV event was jointly genotyped by muCNV (https://github.com/gjun/muCNV), which processes each sample’s CRAM file to generate pileups with all discordant read pairs, split reads, average sequencing depth of each 100-bp interval, average sequencing depth for each candidate event, and sequencing depth curves per GC contents. This per-sample pileup was executed in parallel and then merged into multi-sample pileup files used by muCNV. Joint genotyping was performed based on three different types of information for deletions and duplications: 1) depth-based clustering using Gaussian mixture models. 2) support from discordant read pairs and split reads around the suggested breakpoints, and 3) depth variation of the candidate SV interval relative to its surrounding (+/-1500 bp) genomic region. All sequencing depths were corrected for GC-content using per-sample GC curves calculated during construction of the individual pileup. Inversions calls were made solely based on read pair mapping and split read information, not using sequencing depths. In depth-based clustering, the decision that each consensus event was polymorphic or not was evaluated by Gaussian mixture model estimated by standard expectation-maximization (EM) algorithm by varying number of components. Decision on number of components was based on Bayesian information criterion (BIC). We filtered out variants that had the lowest BIC for a one component model and the variants that had excessive overlap between individual Gaussian components measured by Bhattacharyya coefficients. Deletion and duplication genotypes were also corrected if the SV interval did not show enough depth deviation from its surrounding genomic region. Discordant read-pair and splitread based genotypes for deletions and duplications were called only when it is also supported by sequencing depth profiles. A variant call was made when missing rate is less than 0.5 and at least one variant allele was genotyped.

SVs were annotated using vcfanno, gene and regulatory elements from Ensembl GRCh38, and self-reported ethnicity. We used bcftools filter^24^ to obtain ethnicity-specific information for the subset of common SVs. FST was computed based on Deletions (MAF>0.01) using vcftools^25^

### SV calling on non-human primates

To investigate the potential ancestry allele in non-human primates we downloaded paired end reads from the SRA for the following accessions: ERR2300763, ERR2300767, ERR2300768, SRR490083, SRR490084, SRR490085, SRR490086, SRR490087, SRR490088 and SRR490117. Each of them were mapped to the human genome (GRCh38) using the same pipeline and SVs were called using Parlimanet2 with the same parameters defined previously. The individual call sets were merged using SURVIVOR merge and compared to the full set of the human SV catalog using once a 100bp and once a 1kbp threshold. For the comparison, we did not require that the SV was of the same type, but only focused on agreement of the SV breakpoints.

### ARIC Proteomics Measure

SOMAscan v4 assays (SomaLogic, Inc) were applied to EDTA plasma samples collected from all ARIC visit 5 participants with full consent. SOMAscan uses an aptamer-based array for affinity-proteomic measures^26^. Protein analyte measurements underwent the regular SOMAscan data standardization and normalization process (http://somalogic.com/wp-content/uploads/2017/06/SSM-071-Rev-0-Technical-Note-SOMAscan-Data-Standardization.pdf). Briefly, ARIC samples were median normalized across calibrator samples to remove assay biases within the run. Overall scaling was then performed on a per-plate basis to remove overall intensity differences between runs. Calibration was then performed to remove assay differences between runs followed by median normalization to internal reference. The SOMAscan assay included 5,284 aptamers, and our analysis focused on 4,156 aptamers whose per-plate calibration factors were all within range. The variation of SOMAscan assays were reported elsewhere^27^ with the median coefficient of variation < 5%.

### Statistical Analysis

We investigated aptamer-SV associations in the ARIC study using two scenarios: 1) to use European Americans (n = 3,433) as the discovery samples and replicated the significant findings in African Americans (n = 588); and 2) to use African Americans as the discovery samples and replicated the significant findings in European Americans. Prior to the analysis, we obtained individual aptamer residuals by regressing log2 transformed aptamer values on age, sex, creatinine and study centers, as well as first five ancestry principal components. Individual aptamer residuals were then inverse-normalized to test the linear association with each SV using the GMMAT package^28^ with further adjustment of the same covariates^29^. In scenario 1, significance for discovery was defined as p-value ≤ 1.05e-10 to account for 57,144 SVs with MAC ≥ 5 in European Americans, 4,156 aptamers and two scenarios (0.05/(57144×4156×2)). We brought 663 unique SVs and 403 unique aptamers identified in European Americans to African Americans for replication following the same analytical models. Significance for replication was defined as p-value ≤ 1.87e-7 to account for 663 SVs and 403 aptamers (0.05/663/403). In scenario 2, significance for discovery was defined as p-value ≤ 2.17e-10 to account for 27,655 SVs with MAC ≥ 5 in African Americans, 4,156 aptamers and two scenarios (0.05/(27655×4156×2). We brought 127 unique SVs and 44 unique aptamers identified in African Americans to European American for replication. Significance for replication was defined as p-value ≤ 8.96e-6 to account for 127 SVs and 44 aptamers (0.05/127/44). We applied two-sided Wilcoxon rank-sum test to compare the length of protein-associated deletions and non-protein-associated deletions.

To estimate the impact of SNVs on the SV-protein associations, we related SNVs within ± 1Mb of the identified SVs to protein levels using the same analytical strategies. We compared the effect size on a per-variant basis between genome-wide significant SNVs (defined as p<5e-8) and the identified SV among the participants with both SNV and SV genotypes. In each SV region, we selected tag SNVs if the SNVs reached genome-wide significance and remained significant (defined as p<5e-8) after conditioning on any other SNVs in the same region. We further performed conditional analysis on those 68-protein associations adjusting for tag SNVs. The significance for conditional analyses was defined as SV remained p-value ≤ 1.05e-10 for European Americans discovery set and p-value ≤ 2.17e-10 for African American discovery set.

## Data availability

Structural variation calls are available at dbVar (nstd160) in anonymous form and individual level are available from dbGap (phs001211, phs001395). The list of regions of common SVs clusters are available as supplemental material and https://github.com/orgs/HGSC-NGSI/.

## Acknowledgements

Whole genome sequencing (WGS) for the Trans-Omics in Precision Medicine (TOPMed) program was supported by the National Heart, Lung and Blood Institute (NHLBI). WGS for “NHLBI TOPMed: Atherosclerosis Risk in Communities (ARIC)” (phs001211) was performed at the Baylor College of Medicine Human Genome Sequencing Center (HHSN268201500015C and 3U54HG003273-12S2) and the Broad Institute for MIT and Harvard (3R01HL092577-06S1). Centralized read mapping and genotype calling, along with variant quality metrics and filtering were provided by the TOPMed Informatics Research Center (3R01HL-117626-02S1). Phenotype harmonization, data management, sample-identity QC, and general study coordination, were provided by the TOPMed Data Coordinating Center (3R01HL-120393-02S1). We gratefully acknowledge the studies and participants who provided biological samples and data for TOPMed.

The Genome Sequencing Program (GSP) was funded by the National Human Genome Research Institute (NHGRI), the National Heart, Lung, and Blood Institute (NHLBI), and the National Eye Institute (NEI). The GSP Coordinating Center (U24 HG008956) contributed to cross-program scientific initiatives and provided logistical and general study coordination. The Centers for Common Disease Genomics (CCDG) program was supported by NHGRI and NHLBI, and whole genome sequencing was performed at the Baylor College of Medicine Human Genome Sequencing Center (UM1 HG008898 and R01HL059367).

The Atherosclerosis Risk in Communities study has been funded in whole or in part with Federal funds from the National Heart, Lung, and Blood Institute, National Institutes of Health, Department of Health and Human Services (contract numbers HHSN268201700001I, HHSN268201700002I, HHSN268201700003I, HHSN268201700004I and HHSN268201700005I). The authors thank the staff and participants of the ARIC study for their important contributions.

The Hispanic Community Health Study/Study of Latinos was carried out as a collaborative study supported by contracts from the NHLBI to the University of North Carolina (N01-HC65233), University of Miami (N01-HC65234), Albert Einstein College of Medicine (N01-HC65235), University of Illinois at Chicago (HHSN268201300003I), Northwestern University (N01-HC65236), and San Diego State University (N01-HC65237). The following Institutes/Centers/Offices contribute to the HCHS/SOL through a transfer of funds to the NHLBI: National Institute on Minority Health and Health Disparities, National Institute on Deafness and Other Communication Disorders, National Institute of Dental and Craniofacial Research, National Institute of Diabetes and Digestive and Kidney Diseases, National Institute of Neurological

Disorders and Stroke, NIH Institution-Office of Dietary Supplements. The views expressed in this manuscript are those of the authors and do not necessarily represent the views of the National Heart, Lung, and Blood Institute; the National Institutes of Health; or the U.S. Department of Health and Human Services.

FJS was supported by NIH UM1-HG008898. BY was in part supported by NIH R01HL142003.

